# Advancing Wildlife Image Analysis: A Graph Attention Contrastive Learning Approach for Region-Specific Mammal Classification

**DOI:** 10.1101/2025.09.17.676694

**Authors:** Youngmin Kim, Cheol-Han Kim, Chang-Seob Yun, Gea-Jae Joo

## Abstract

1. Camera traps have become a cornerstone of wildlife ecological research, yet the manual analysis of the millions of images they generate requires substantial time and resources. Deep learning-based automation has emerged as a promising solution, existing global general-purpose models exhibit limitations in precisely recognizing local endemic species and adapting to unique local ecosystems.

2. This study developed a high-performance classification model optimized for native species. A large-scale “Korean Wildlife Dataset” was constructed from data collected across diverse domestic habitats, and a novel architecture was proposed to overcome limitations of conventional CNNs. The proposed Graph Attention Contrastive Learning (GACL) model is structured as a two-stage pipeline. Stage one employs YOLOv5 and MegaDetector to detect animals, humans, and vehicles, filtering valid images. Stage two performs fine-grained species classification. GACL captures structural relationships among object parts using a Graph Attention Transformer (GAT) and aligns semantic correspondence between images and textual descriptions via Parallel Contrastive Learning, enabling deeper understanding beyond simple visual features.

3. Evaluation on an independent test set demonstrated that the proposed model robust classification performance with an overall accuracy of 96.83% across four classes (Wildboar, Goral, Deers, and Other). Notably, in a comparative analysis against a global general-purpose model, our model showed distinct advantages in the precise recognition of endemic species. Furthermore, it exhibited a lower false positive rate in identifying animals in empty images, confirming its potential to enhance the efficiency of the data cleaning process.

4. Beyond technical accuracy, this study highlights that ’region-specific AI models’ that reflect local ecological characteristics can provide substantial practical value for wildlife monitoring and biodiversity conservation. Future work will require continuous efforts in data diversification and model lightweighting to further improve model robustness and practicality.

## 1. Introduction

The use of camera traps has become a cornerstone of modern ecological research, enabling continuous and non-invasive monitoring of wildlife across vast and often inaccessible environments (Burton et al., 2015). Recent technological improvements, including enhanced hardware durability and extended battery life, have further strengthened their utility, allowing reliable data collection even under extreme climatic conditions (Swanson et al., 2015). However, these advancements have also introduced a significant challenge: camera trap deployments generate massive image datasets, often comprising millions of frames. Manual processing of these data represents a major bottleneck, consuming substantial human resources and limiting the scalability of ecological studies (Norouzzadeh et al., 2018). To select valid data for precise wildlife population studies, it is essential not only to detect the mere presence of wildlife but also to extract accurate species identifications, the extraction of metadata such as timestamp, location, and temperature, and, in some cases, the analysis of finer-grained details including group size, behavioral states, or event duration (Swanson et al., 2015; Tabak et al., 2019). The interpretation of such qualitative information requires substantial expertise and considerable analytical effort. When conducted manually, this process is prone to errors stemming from operator proficiency and fatigue, which may ultimately compromise the reliability and efficiency of the overall data analysis (Norouzzadeh et al., 2018).

Despite these advances, critical limitations remain. First, general-purpose models are often trained on datasets from geographically restricted regions, such as North America or Africa, and frequently struggle to accurately identify endemic species in ecologically distinct areas. For instance, key species of the Korean Peninsula, including Water deer (*Hydropotes inermis*), Siberian chipmunk (*Tamias sibiricus asiaticus*), and Raccoon dog (*Nyctereutes procyonoides*), are poorly recognized by models trained on foreign fauna. Second, conventional CNN architectures focus primarily on object appearance, without explicitly modeling the structural relationships among object parts or incorporating rich semantic context from associated metadata— information that is critical for distinguishing between morphologically similar species in complex natural environments. These limitations underscore the need for both region-specific datasets and novel architectures that can capture higher-order structural and semantic information.

This study confronts these challenges directly. We first constructed a large-scale, regionally tailored “Korean wildlife dataset” based on long-term camera trap collections. Leveraging this dataset, we developed Graph Attention Contrastive Learning (GACL), a novel deep learning architecture designed to overcome the limitations of existing models by integrating structural and semantic cues. Through rigorous evaluation, we demonstrate the enhanced accuracy of our approach for local species and highlight the critical importance of region-specific, architecture-aware solutions for ecological monitoring. Ultimately, this work establishes a technical foundation for more reliable and efficient intelligent systems in biodiversity conservation management.

## 2. Theoretical Background and Related Work

### 2.1 Deep Learning-Based Image Recognition for Wildlife Monitoring

mage-based monitoring technologies for wildlife conservation and ecosystem management have steadily advanced over the past decades. However, data collected by automated sensors, such as camera traps, often contain complex backgrounds and non-uniform lighting conditions, making it difficult for traditional image processing techniques to achieve high analytical accuracy. The advent of deep learning has marked a pivotal shift in overcoming these challenges, as it enables predictive modeling by autonomously learning visual patterns directly from image data (A Krizhevsky et al., 2012). Through this hierarchical feature learning, CNNs progress from recognizing low-level features, such as edges, shapes, and textures, to high-level structural and morphological patterns. Such hierarchical feature learning not only supports single-object classification but also facilitates more complex visual tasks, including object detection, image segmentation, and behavior recognition, thereby demonstrating strong applicability to ecological image analysis (Z Zou et al., 2023; A Mathis et al., 2018).

The core objective of wildlife monitoring is to accurately detect animals in images and classify them at the species level. To this end, state-of-the-art object detection algorithms such as YOLO, Faster R-CNN, and RetinaNet have been developed, all of which are built upon CNN architectures. These models typically leverage pre-trained backbones, including ResNet and EfficientNet, to extract hierarchical features from images and determine not only ‘*which’* animal is present but also ‘*where’* it is located (Bochkovskiy et al., 2020; Liu et al., 2016). By incorporating both low-level and high-level morphological traits—such as ears, tails, and body shapes—these models enable rapid and accurate extraction of meaningful information from large-scale datasets, maintaining high performance even under variable backgrounds, illumination conditions, and inter-species morphological similarities.

### 2.2 CNN-Based Tools for Wildlife Image Analysis

The challenge of efficiently analyzing vast camera trap datasets has been largely addressed by the development of specialized CNN-based tools, with YOLO and MegaDetector emerging as two of the most influential.

YOLO (You Only Look Once) revolutionized real-time object detection by introducing a ’one-shot’ approach that simultaneously detects and classifies objects within a single pass of a CNN (Redmon et al., 2016). Its exceptional processing speed makes it highly suitable for the rapid analysis of large-scale ecological datasets. Moreover, YOLO has the ability to interpret objects within the global context of the entire image, integrating environmental cues such as background and spatial structure instead of relying solely on localized features. This capability has proven particularly valuable for distinguishing between morphologically similar species in complex natural backgrounds. These structural advantages have led to its widespread application in ecological monitoring, where it enables automated, high-volume processing of camera trap data with both efficiency and reliability (Ma Zhibin et al., 2024).

Building on this foundation, MegaDetector was developed by Microsoft’s AI for Earth project as an open-source tool specifically optimized for the initial filtering of camera trap data. It leverages pre-trained CNN backbones (such as ResNet and EfficientNet) trained on an extensive dataset of camera trap images to automatically detect objects (Beery et al., 2019; Microsoft, 2020). Instead of performing fine-grained species classification, MegaDetector focuses on three high-level classes—animals, people, and vehicles—effectively separating images containing wildlife from empty frames and substantially reducing the manual labor required for data review (Fennell M et al., 2022). By providing this foundational step, MegaDetector maximizes the efficiency of subsequent tasks, including species identification and behavior analysis.

### 2.4 Related Work and Limitations

The advancement of deep learning-based object detection has extended beyond general-purpose tools like MegaDetector, leading to the development of large-scale application platforms for analyzing and sharing field data. A prominent example is **Wildlife Insights** (www.wildlifeinsights.org/), which integrates camera trap data from researchers worldwide and provides AI-analyzed results, making a substantial contribution to global biodiversity conservation efforts. Such platforms relieve individual researchers of data processing burdens, enabling them to focus more fully on ecological questions (Ahumada et al., 2020). However, these global platforms and general-purpose models have clear limitations. First, their training data are heavily biased towards the fauna of specific regions, such as North America and Africa, which can lead to relatively degraded recognition performance for key endemic species of the Korean Peninsula, including the water deer (*Hydropotes inermis*), the leopard cat (*Prionailurus bengalensis*), and the raccoon dog (*Nyctereutes procyonoides*). Second, while models such as MegaDetector excel at detecting objects within broad categories, such as “animals,” they are insufficient for fine-grained species-level classification without additional downstream models and region-specific training datasets. In conclusion, the successful application of these robust tools to local ecological monitoring requires a foundational study to first develop and validate models trained on datasets tailored to the local environment and species composition.”

## 3. Materials and Method

In this study, we aimed to improve the automatic detection and classification of wildlife native to Korea by developing a deep learning-based image analysis model that accounts for the unique characteristics of the local ecological environment. To this end, we systematically curated a high-quality training dataset and then developed and trained a custom two-stage architecture for detection and classification.

### 3.1. Data Collection and Training Dataset Construction

The performance and generalization capability of deep learning models are largely determined by the quantity and quality of training data. In this study, we prioritized the construction of a high-quality dataset that captures the diversity of the local environment. The dataset construction process was implemented through a structured pipeline, beginning with the planning stage, followed by the acquisition of raw images, data cleaning and preprocessing, meticulous annotation, augmentation, and culminating in the compilation of the final training dataset.

During the initial stage of data collection and class definition, training data were compiled from images captured over several months to years using camera traps across diverse habitats in Korea. The classification scheme was defined with four classes: ‘Wildboar’ (*Sus scrofa*), ‘Goral’ (long-tailed goral, *Naemorhedus caudatus*), ‘Deers’ (including roe deer (*Capreolus pygargus*) and water deer (*Hydropotes inermis*)), and an ‘Other’ comprising additional mammals and birds. Data collection aimed to secure a total of 60,000 images, with 15,000 images allocated to each class. Images were carefully curated to ensure sufficient diversity and representativeness in terms of morphological traits, behavioral patterns, and varying capture conditions for each category. In the data refinement and quality control phase, images deemed unsuitable for model training or likely to degrade model performance were systematically removed. Specifically, low-resolution or blurry images in which essential features for species identification, such as body contours or fur texture, were obscured were excluded. This included cases affected by camera shake, excessive noise during nighttime capture, or images in which animals were partially hidden by vegetation, captured from a distance as silhouettes, or where only a small portion of the animal was visible. To further enhance dataset quality and reduce redundancy, sequences capturing the same individual in similar poses at the same location were pruned, retaining only representative frames. Additionally, to mitigate potential overfitting arising from class-specific perspective biases (e.g., frontal or lateral views), images were sampled to balance the dataset and include a variety of viewing angles (Fig 1).

**Figure 1.**
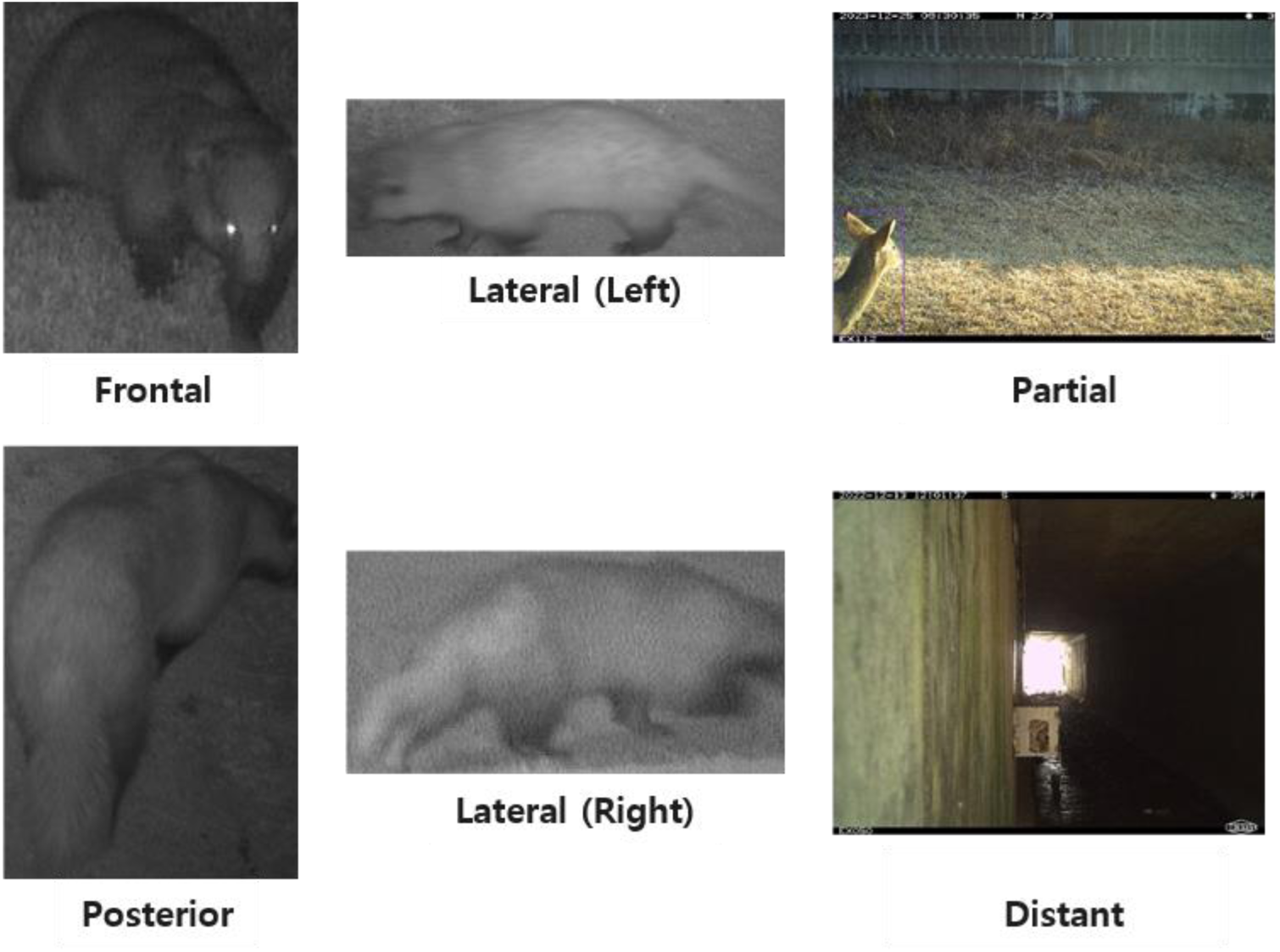
Examples of Image types curated for the training dataset. The dataset was curated to include diverse perspectives (frontal, posterior, lateral, partial and distant views) Following data refinement, a multi-faceted data augmentation strategy was employed to expand the dataset and enhance model robustness. Data augmentation is a key regularization technique that artificially generates new training samples from a limited set of original images, thereby preventing the model from overfitting to the training data and ensuring robust generalization to diverse real-world variables (Hernández-García, A., & König, P. 2018; Krizhevsky, A. et al., 2017). In this study, we applied a suite of representative augmentation techniques included geometric transformations (resizing for scale invariance and random rotation for orientation invariance), photometric transformations (adjustments to brightness, exposure, hue, and chroma using both HSV and LAB color spaces), and the advanced regularization technique Mixup (Zhang et al., 2018). For photometric transformations, we adopted a balanced, dual-space approach to chroma augmentation leveraging both HSV and LAB color spaces. This allowed us to simulate diverse environmental conditions, with HSV transformations producing more dramatic variations in hue, saturation, and value channels, while LAB transformations provided perceptually natural shifts along the green–red (a*) and blue–yellow (b*) axes. The effectiveness of these color-based augmentations is summarized in Figure 2, which illustrates how the overall color distribution broadened and the Shannon entropy increased, indicating improved robustness to varied illumination and color conditions. Detailed per-channel statistics and quantitative metrics, including KL divergence, are provided in Appendix A.

**Figure 2.**
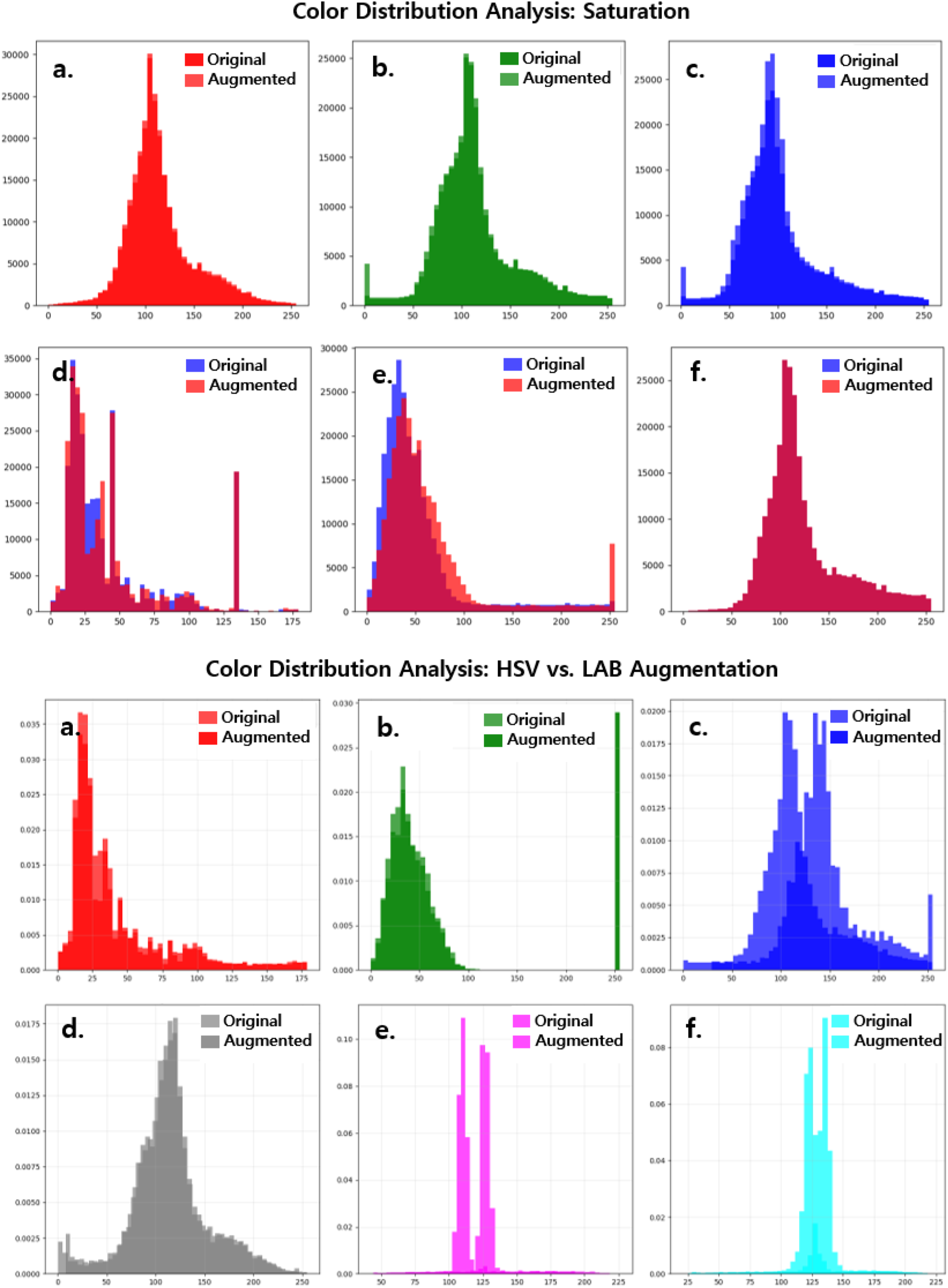
Quantitative analysis of color distribution for photometric augmentations. Top: Histograms showing the effect of brightness augmentation on RGB (a-c) and HSV (d-f) channels. Bottom: Histograms comparing the effects of chroma augmentation in HSV (a-c) and LAB (d-f) color spaces.

The Mixup technique was further applied to enhance model generalization by linearly interpolating between two distinct images and their labels, encouraging smoother decision boundaries. Figure 3 visualizes the mixing ratio (λ) sampled from a Beta distribution, showing that approximately 42% of the samples retain over 95% of one original image’s features. This ensures that synthetic samples largely preserve the distinctive traits of their source classes while also exploring intermediate feature combinations, contributing to improved robustness across unseen data.

**Figure 3.**
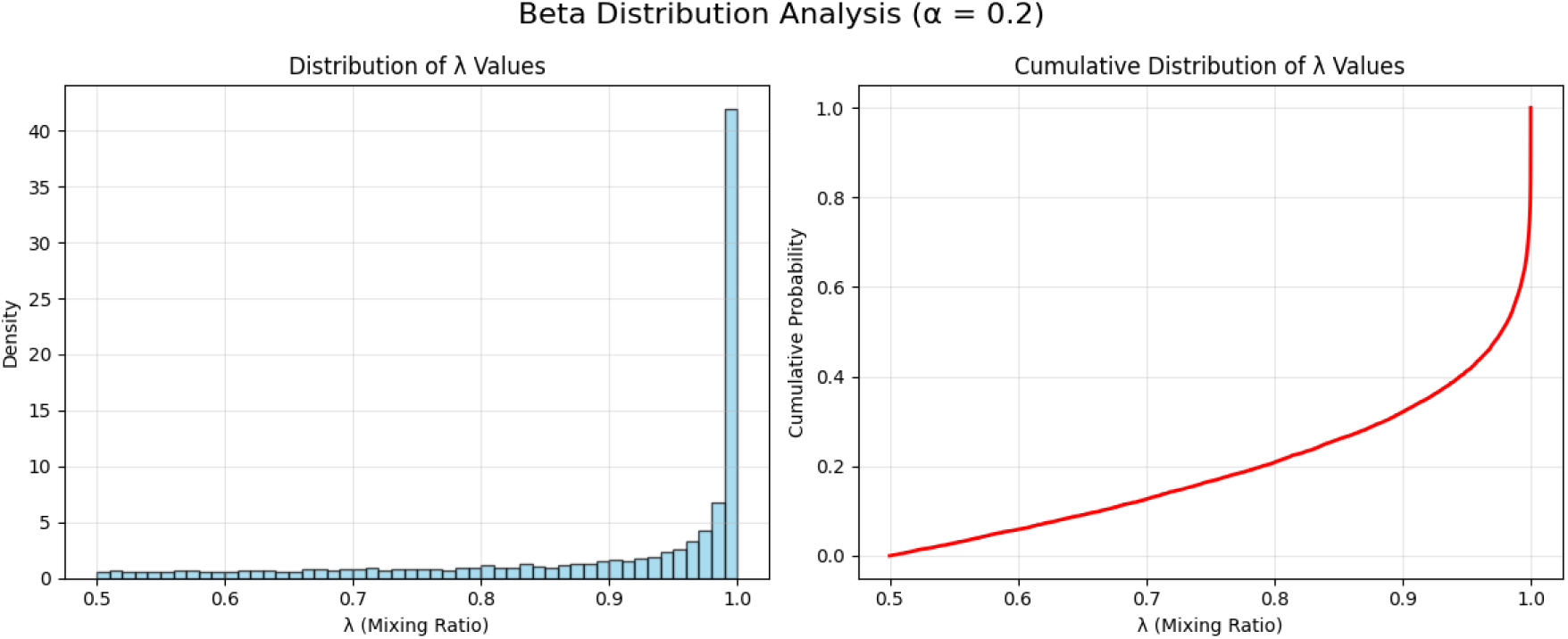
Beta distribution analysis of the mixing ratio λ. Probability density function (left) and cumulative distribution function (right).

### 3.2 Model Architecture

The automated analysis system proposed in this study consists of a sequential pipeline with two stages: (1) wildlife object detection using a general-purpose model, and (2) fine-grained species classification.

In the first stage, the object detection model aims to rapidly identify valid images containing wildlife from millions of raw images, thereby reducing the computational burden for subsequent processing. For this purpose, model training was conducted using MegaDetector and YOLOv5. The classification scheme for this stage was limited to three high-level classes, ’animal’, ’person’, and ’vehicle’, to efficiently separate images containing animals from non-animal images. Only images detected as ’animal’, along with their bounding box information, are passed to the next stage for species classification. In the second stage, the species classification model utilizes a Graph Attention Contrastive Learning (GACL) approach to address the limitations of conventional CNN-based methods. Whereas traditional models focus solely on the visual features of individual objects, our approach simultaneously captures the relationships among object parts and the semantic correspondence between images and text. This dual focus enhances classification accuracy. The model consists of three core components: a GAT encoder, a text encoder, and a parallel contrastive learning framework.

The first core component, the GAT (Graph Attention Transformer) encoder, is responsible for encoding both the visual and structural information of the input image. This process proceeds sequentially through object detection, patch formation, positional embedding, and graph construction (Fig 5). Initially, a YOLOv5 model identifies the detailed parts of the animal, and a tracking model groups patches based on each object’s tracking ID. Each patch, representing a characteristic region of the object, is passed through a convolutional neural network with multiple dilation rates to extract features. This multi-dilated convolutional network enables the GAT encoder to simultaneously capture information across various spatial scales (Table 1).

**Figure 4.**
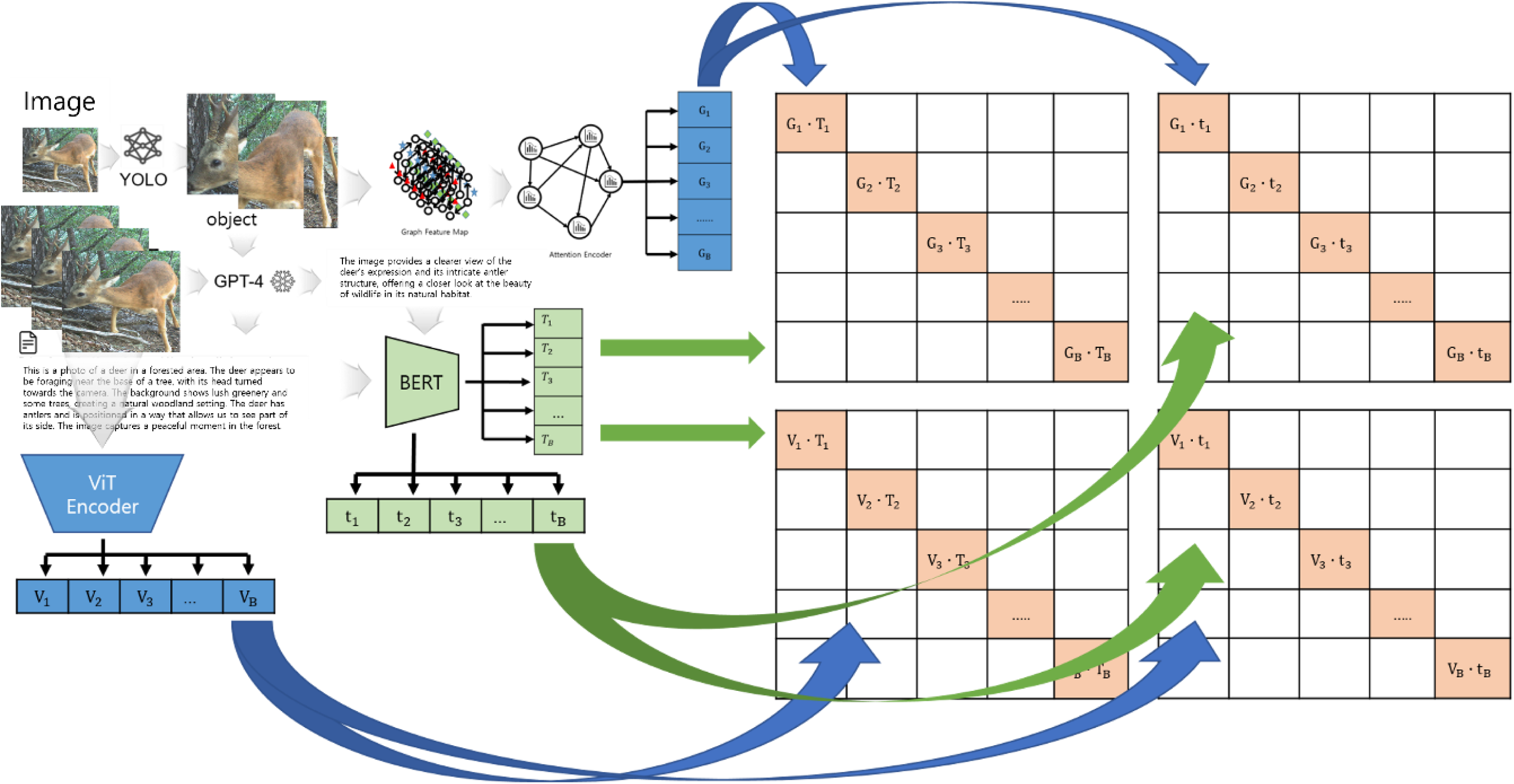
Architecture of the GACL Model. The proposed Graph Attention Contrastive Learning (GACL) model processes an input image and a corresponding text description through three parallel encoders. An Attention Encoder (GAT) extracts global structural features (G) from a graph representation of the detected object. Concurrently, a ViT Encoder extracts local patch features (V) from the image, and a BERT Encoder extracts both global (T) and local (t) semantic features from the text. These four feature types are then aligned using Parallel Contrastive Learning, which simultaneously compares them across four relationships: global-to-global (G-T), global-to-local (G-t), local-to-global (V-T), and local-to-local (V-t).

**Figure 5.**
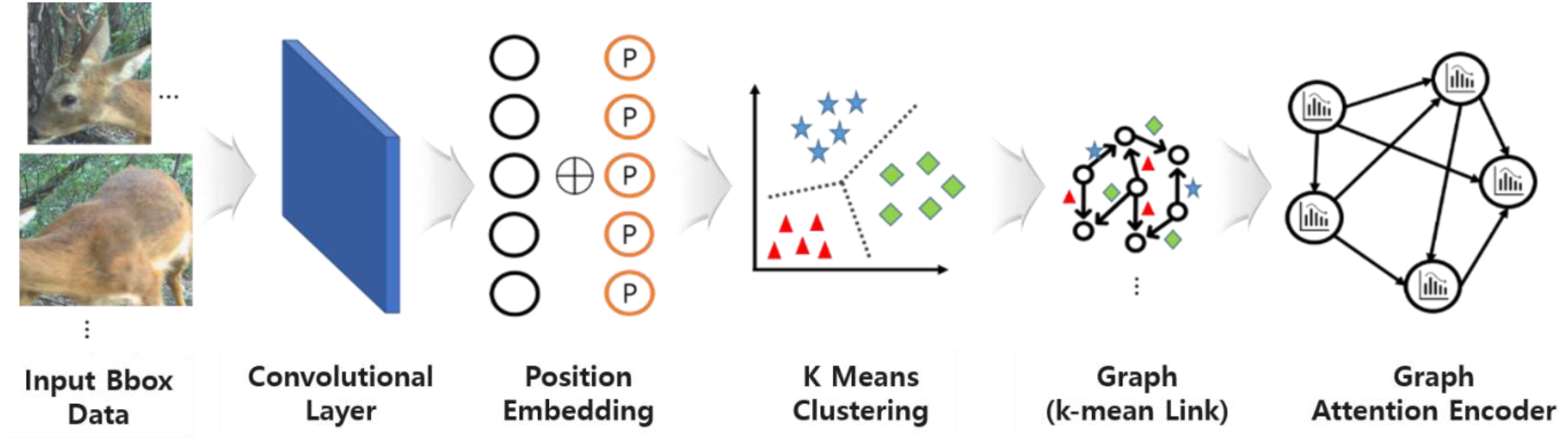
The generation process of input data for the GAT encoder.

**Table 1.** Multi-Dilated Convolutional Network Architecture.

|  | Kernel | Dilation Rates | Feature Description |
| --- | --- | --- | --- |
| Conv2D | 3x3 | dilation=1 | Captures fine-grained local features (similar to standard CNN filters) |
| Conv2D |  | dilation=2 | Collects information from a broader local region |
| Conv2D |  | dilation=4 | Detects larger structural patterns of the object |
| Conv2D |  | dilation=5 | Learns global information by considering an expanded receptive field |

Specifically, multiple Conv2D layers with 3×3 kernels but varying dilation rates operate in parallel. Standard convolutions (dilation rate = 1) focus on fine-grained local features such as fur texture or eye shape. As the dilation rate increases (2, 4, 5), the network captures progressively broader structural patterns and global context. By considering multiple receptive fields simultaneously, the model captures both detailed local features and overall contextual information. Positional embeddings preserve the spatial relationships of each patch, maintaining relational context throughout processing.

Graph construction follows to represent relationships among object patches. Nodes correspond to patches, while edges are defined via K-means clustering based on visual similarity. Nodes within the same cluster receive strong weights, whereas inter-cluster edges are weighted weakly, forming a graph that reflects spatial patterns among object parts. This graph serves as the input to the GAT-based encoder. As depicted in Figure 6, the GAT encoder is structured as a series of repeating blocks. Each block contains a GATv2Conv layer, Layer Normalization, and a Multi-Layer Perceptron (MLP), with residual connections encompassing the operations. The GATv2Conv layer dynamically attends to neighboring nodes, selectively gathering and updating features. Layer Normalization stabilizes the data distribution, and the MLP transforms these features into a more complex, high-dimensional representation. Residual connections mitigate potential information loss, ensuring that each node’s original spatial characteristics are preserved while relational features are learned. This architecture allows the GAT encoder to effectively emphasize features critical for species classification (e.g., the antlers of a roe deer) and generate rich feature vectors that encapsulate both the overall structure and detailed components of each object.

**Figure 6.**
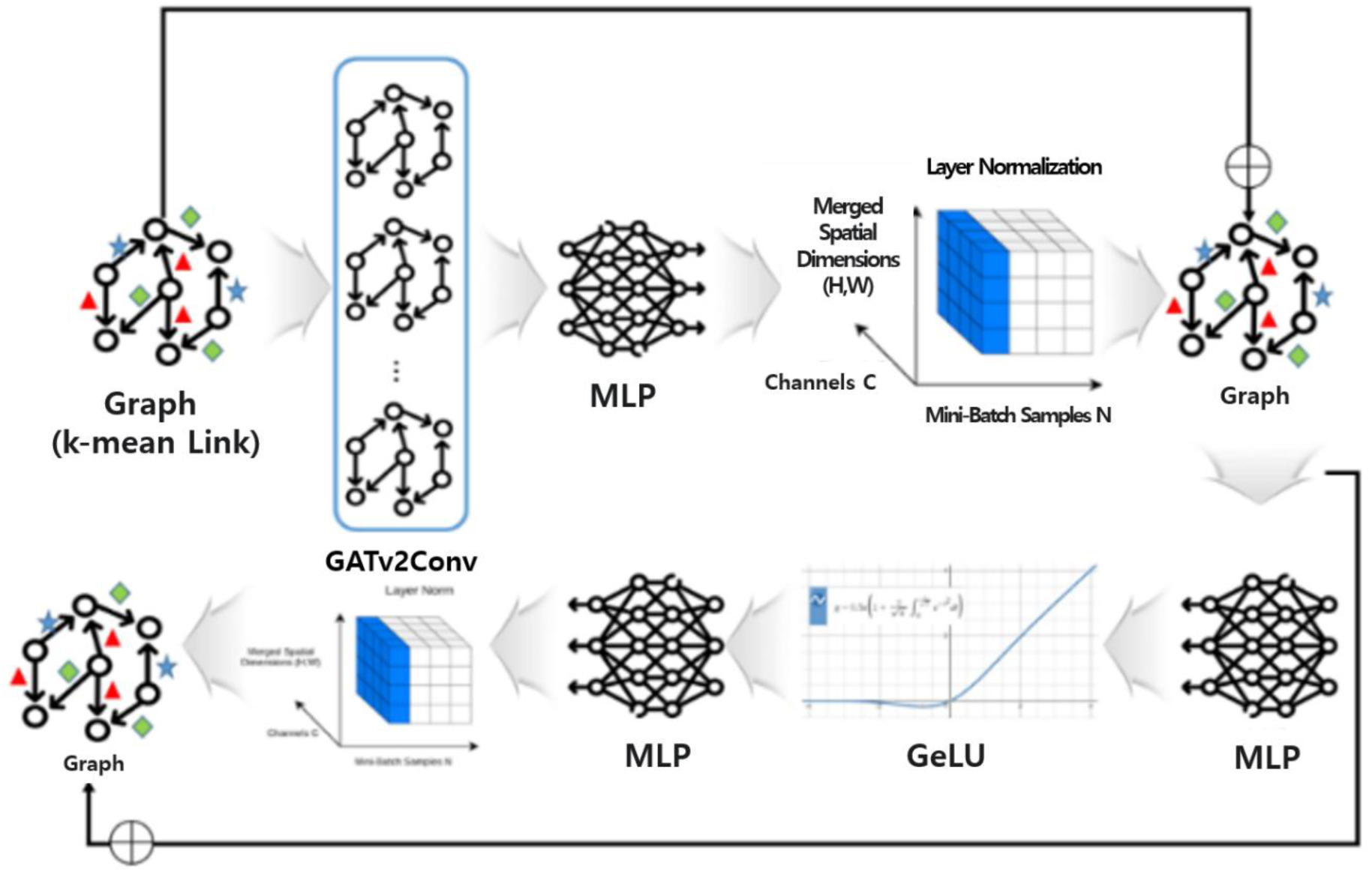
Detailed Architecture of the GAT Encoder. The input graph is processed through a series of blocks, each applying graph attention (GATv2Conv), normalization, and non-linear transformation (MLP) with a residual connection.

The text encoder, the second core component, provides semantic information about each object. It encodes natural language prompts using a BERT (Bidirectional Encoder Representations from Transformers)-based model, capturing meaningful context from textual descriptions. Two types of prompts are employed (Figure 7):

**Figure 7.**
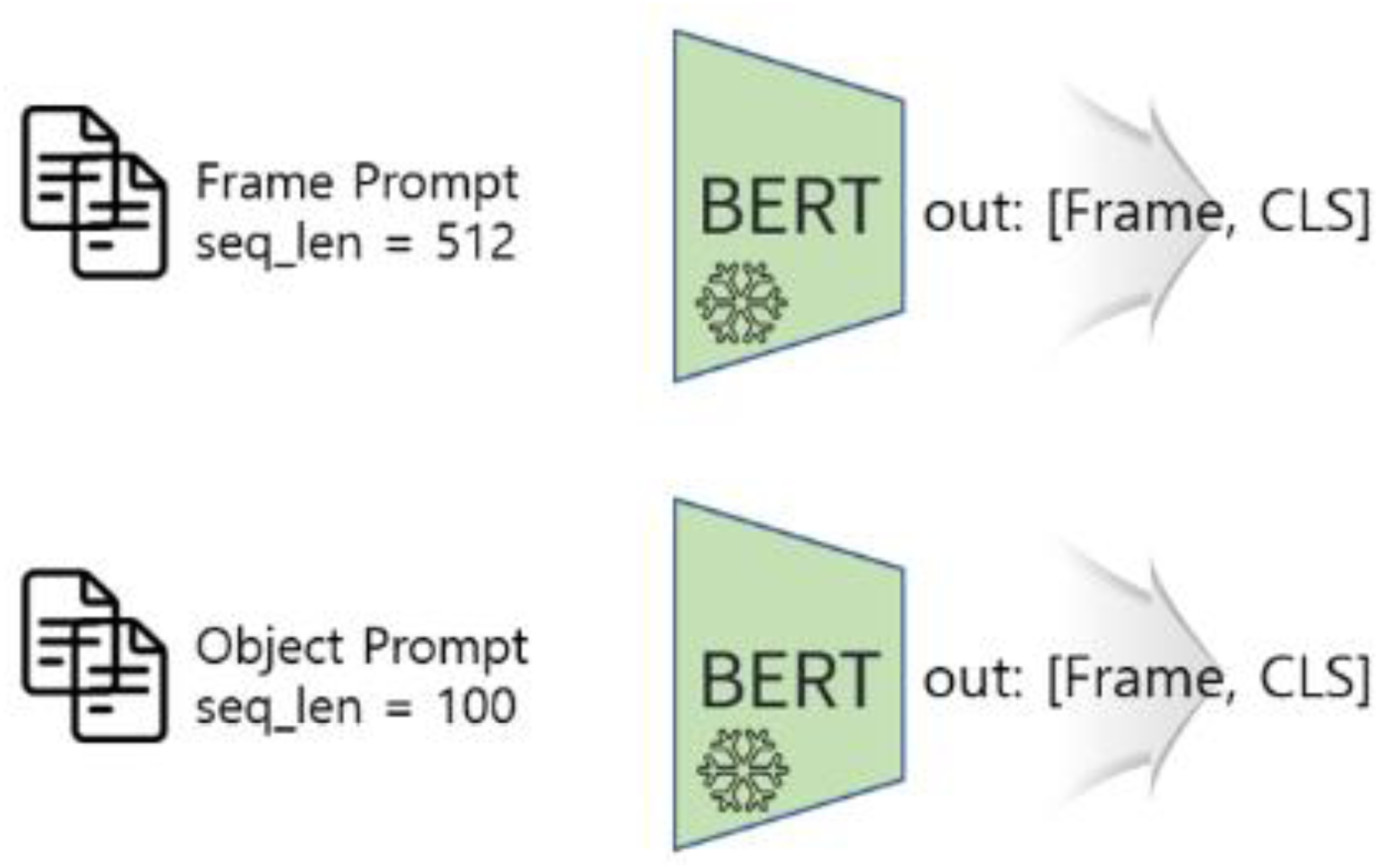
The text encoding process. The BERT encoder processes two types of prompts with different maximum sequence lengths to generate semantic feature vectors.

### Frame Prompt

Provides detailed, long descriptions of the entire object or its context, processed up to a maximum length of 512 tokens (seq_len = 512).

### Object Prompt

Provides concise, focused descriptions of specific parts of the animal (e.g., antlers or legs), processed up to a maximum length of 100 tokens (seq_len = 100).

Each prompt is generated based on a large language model GPT-4, and the pre-trained BERT model utilizes its self-attention mechanism to extract semantically significant words and contextual relationships. Both prompt types, despite their differing lengths and detail, are transformed into fixed-dimensional semantic feature vectors. These vectors are subsequently used in the contrastive learning stage to align with corresponding image feature vectors.

The third and final core component is the Parallel Contrastive Learning process, which serves as the central learning mechanism of the proposed model. In this stage, image feature vectors (*v*_*j*_i) from the GAT encoder and text feature vectors (*w*_*j*_) from the BERT-based text encoder are aligned within a shared embedding space using a contrastive learning approach.

The fundamental mathematical principle of this contrastive learning is illustrated in Figure 8. For each image-text pair, a cosine similarity *S*_(*i*,*j*)_is computed and scaled by a temperature parameter (*τ*)), with higher scores indicating greater semantic proximity. These scores are converted into probabilities via a Softmax function, representing the likelihood of correct image-text correspondence in both directions (Pⱼ|i) and (Pi|ⱼ). The model is trained to minimize a contrastive loss, averaging the image-to-text loss(L_iₘₖ_→_ₜₓₜ_) and text-to-image loss (L_ₜₓₜ_→_iₘₖ_) to optimize bidirectional semantic alignment.

**Figure 8.**
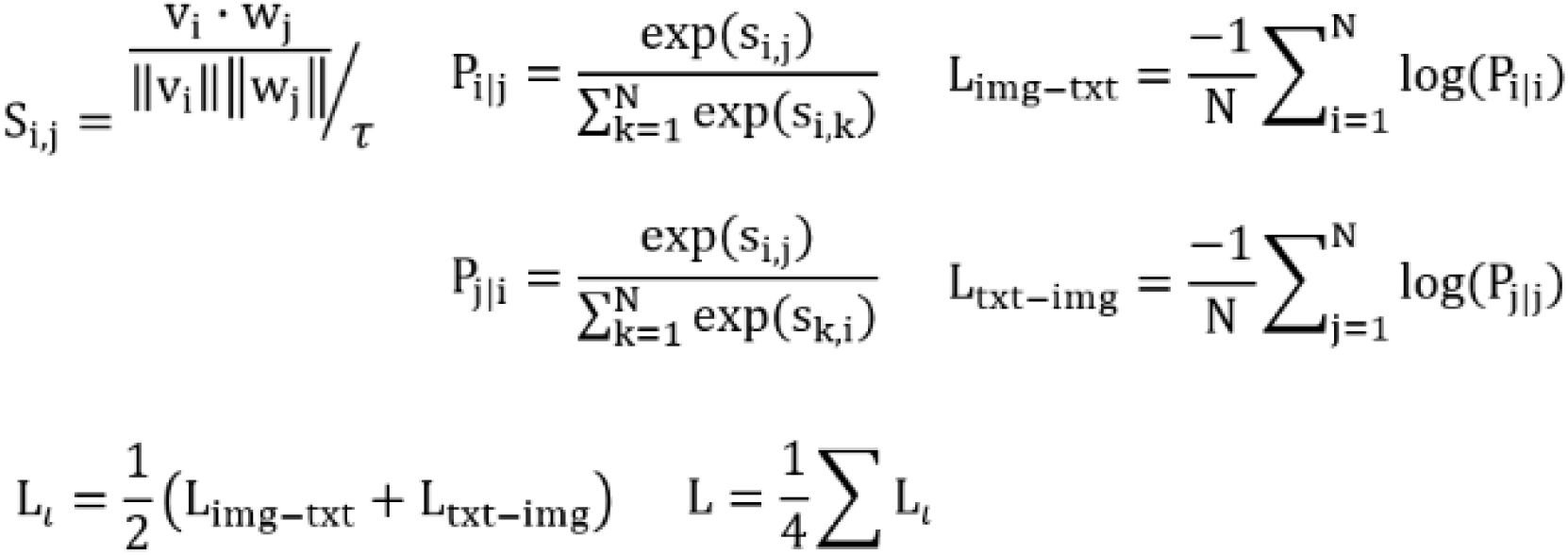
The mathematical principle of Bidirectional Contrastive Learning.

Unlike conventional unidirectional methods, our model extends this bidirectional contrastive learning across four parallel levels of semantic relationships (Fig 9):

1. **Local-to-Local:** Aligns partial image features (e.g., antlers) with corresponding partial text features (e.g., “pointed antlers”) to capture fine-grained details.
2. **Local-to-Global:** Aligns partial image features with full text descriptions (e.g., “a brown animal with antlers”) to contextualize local cues.
3. **Global-to-Local:** Aligns full image features with partial text to identify local details within the global context.
4. **Global-to-Global:** Aligns full image features with full text descriptions for overall contextual consistency.

**Figure 9.**
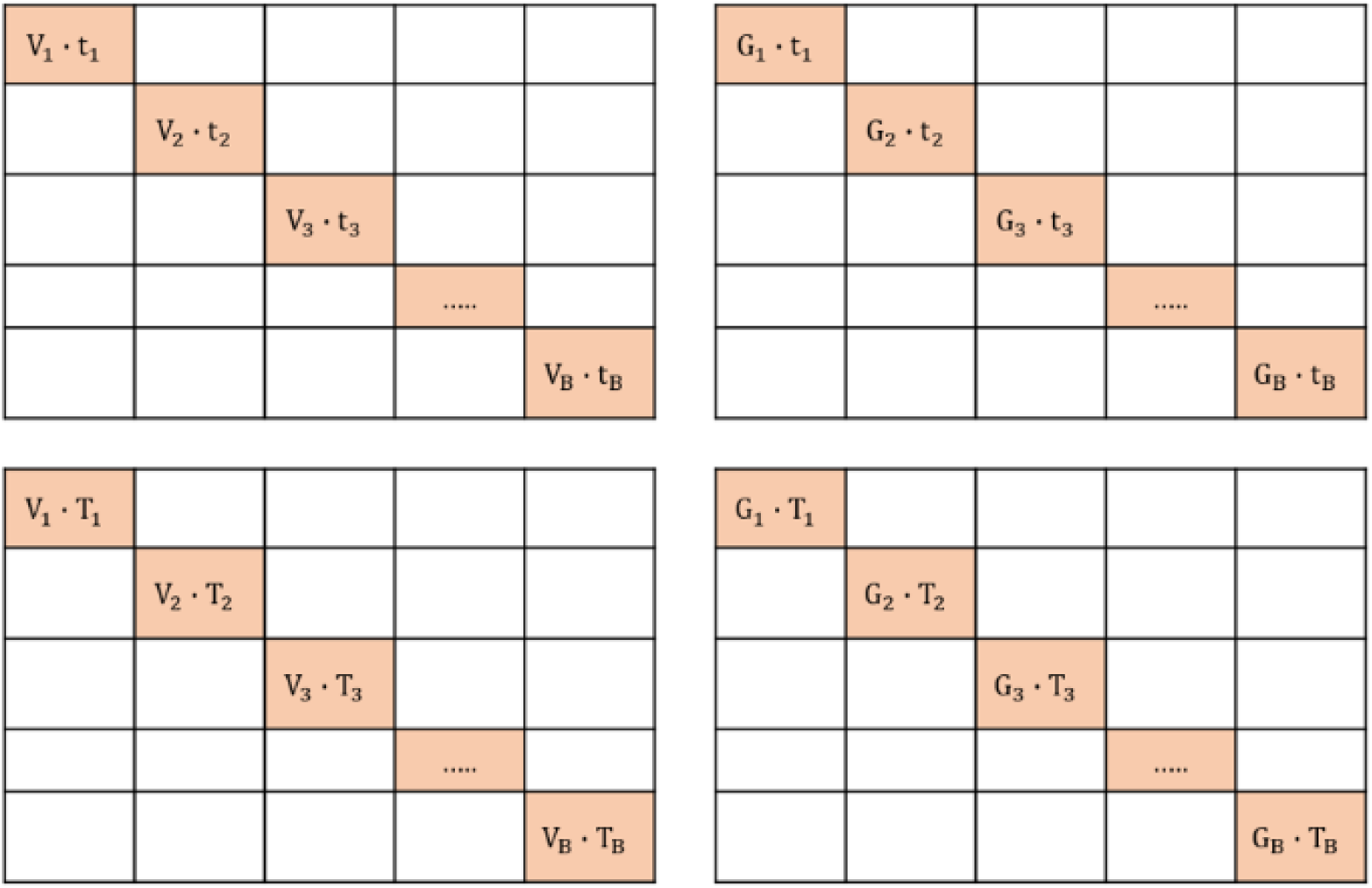
The Parallel Contrastive Loss components. The loss is computed in parallel across four relationships: local-to-local (V-t), global-to-local (G-t), local-to-global (V-T), and global-to-global (G-T). (where V: local image, G: global image, t: local text, T: global text).

The total loss is defined as the average of these four components, ensuring that local and global semantic relationships are jointly optimized. This parallel strategy allows errors at one level to be compensated by others, resulting in more robust and precise semantic alignment and ultimately maximizing classification performance.

In conclusion, the proposed Graph Attention Contrastive Learning (GACL) model integrates the Graph Attention Transformer (GAT) with Parallel Contrastive Learning to offer a novel approach for wildlife image analysis. While conventional CNN-based classification models focus primarily on learning individual object features and struggle to capture interactions and contextual relationships, the proposed model overcomes this limitation by representing and learning inter-object relationships in a graph structure through the GAT framework. This approach is particularly advantageous for high-dimensional classification tasks. Furthermore, by employing a Parallel Contrastive Loss to align multi-modal data, the model achieves more precise classification and retrieval capabilities compared to traditional single-object classification methods.

### 3.3 Processing Pipeline

The detection and classification models are integrated into a single, comprehensive processing pipeline, as illustrated in Figure 10. This pipeline defines the full workflow, from raw image input to final analytical results, in which sequentially operating modules perform distinct, specialized functions.

**Figure 10.**
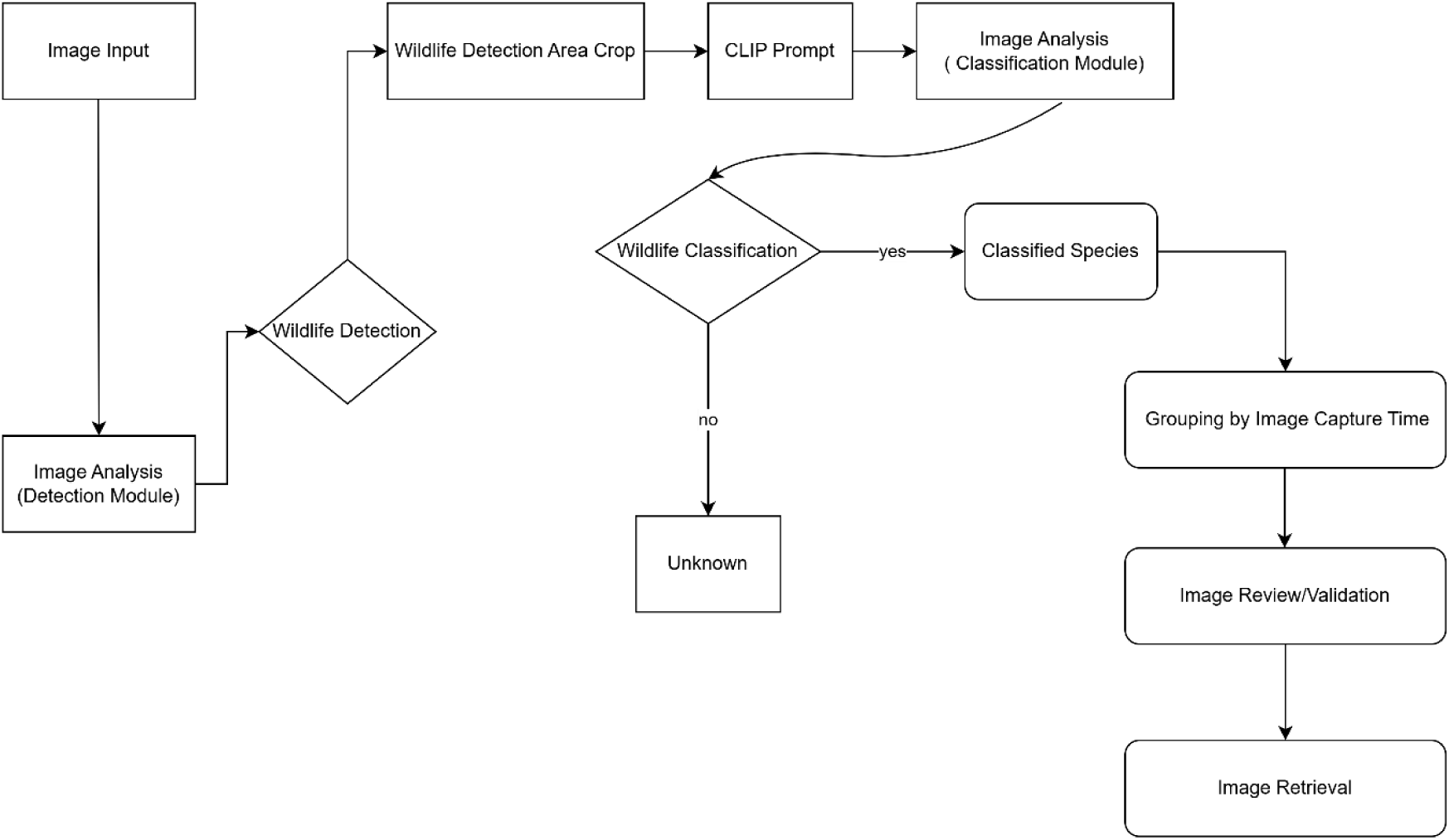
Flowchart of the overall processing pipeline.

In the initial stage, raw images are fed into the Stage 1 Detection Module, which rapidly determines the presence of wildlife. If an animal is detected (Wildlife Detection: yes), the corresponding object region (Bounding Box) is precisely extracted (Wildlife Detection Area Crop). This cropped region undergoes preprocessing, including CLIP prompt processing, to enhance the performance of the subsequent Stage 2 Classification Module. The Classification Module employs the proposed Graph Attention Contrastive Learning (GACL) framework to perform fine-grained species classification (Wildlife Classification). The model is intentionally designed to minimize classification errors: it either assigns the object to a pre-trained class or, if classification confidence falls below a predefined threshold, categorizes it as an unidentified species. Classified results (Classified Species) are not limited to individual image labels. In post-processing, images are grouped by capture time (Grouping by Image Capture Time) to consolidate temporally correlated images into single events. Finally, these AI-processed results undergo a review stage (Image Review/Validation), allowing researchers to verify, approve, or correct them before storage in a retrievable database.

## 4. Results: Performance Evaluation

### 4.1 Evaluation setup and Metrics

To objectively and quantitatively evaluate the performance of the proposed deep learning model, four target classes were defined: Wildboar, Goral, Deers, and Other species. The test set was curated from 11,065 independent images that were not used during training, thereby reflecting the characteristics of real-world field data. This set was constructed as an imbalanced dataset, consisting of 984 Wildboar, 1,979 Deers, 1,108 Gorals, and 6,994 images of Other species.

The quantitative performance of the model was measured from multiple perspectives using standard evaluation metrics in the fields of classification and object detection. The fundamental classification outcomes are defined as follows:.

**True Positive (TP):** the model correctly predicts a sample as positive when it is actually true. **False Positive (FP):** the model incorrectly predicts a sample as positive when it is actually false (Type I error).

**True Negative (TN):** the model correctly predicts a sample as negative when it is actually false. **False Negative (FN):** the model incorrectly predicts a sample as negative when it is actually true (Type II error).

Based on these outcomes, precision and recall were employed as primary metrics to indicate the correctness of the model’s predictions. Precision 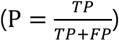 measures the proportion of samples predicted as a given class that are actually correct, and is closely related to false positives. Recall 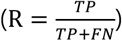 measures the proportion of actual samples of a class that were correctly identified, and is closely related to false negatives. Since precision and recall inherently exhibit a trade-off relationship, their harmonic mean, the F1-Score 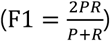, was employed to assess balanced classification performance.

For object detection, the Mean Average Precision (mAP) was used as a comprehensive indicator of both classification accuracy and localization precision. mAP is derived by calculating the Average Precision (AP), defined as the area under the precision–recall curve for each class, and then averaging across all classes. Given its robustness to class imbalance, mAP is particularly suitable for datasets such as ours, which are both large-scale and highly imbalanced.

Finally, to qualitatively analyze misclassification patterns, a Confusion Matrix was employed. This visualization enables not only an assessment of per-class classification accuracy but also an intuitive understanding of systematic misclassification trends.

### 4.2. Model Performance and Analysis

The evaluation using the independent test dataset demonstrated that the proposed model achieved an overall classification accuracy of 96.83%, indicating strong performance. A class-wise analysis further confirmed stable classification ability, with all classes achieving an F1-score above 0.94 (Table 3). Notably, the “Other” and “Deers” classes reached F1-scores of 0.99 and 0.97, respectively, suggesting that their morphological features were effectively captured by the model. When assessed using mean Average Precision at IoU threshold 0.5 (mAP@0.5), a key metric to comprehensively evaluate both detection and classification accuracy, the model consistently performed well, with scores above 0.81 across all classes. In particular, the Other class achieved the highest mAP@0.5 score of 0.90, highlighting the model’s ability to accurately localize and identify diverse animal types within this category.

**Table 2.** Key Hyperparameters of the BERT Model.

| Hyperparameter | Description |
| --- | --- |
| L=12 | Number of Transformer encoder layers (stacked self-attention layers) |
| D=768 | Hidden size (dimensionality of embeddings and encoder outputs) |
| A=12 | Number of attention heads in the Multi-Head Self-Attention mechanism |
| 110M | Total number of model parameters (110 million) |

**Table 3.** F1-Score and mAP results for the proposed model.

| Class | Precision | Recall | F1-score | mAP |
| --- | --- | --- | --- | --- |
| Wildboar | 0.91 | 0.96 | 0.94 | 0.81 |
| Goral | 0.96 | 0.96 | 0.96 | 0.89 |
| Deers | 0.97 | 0.97 | 0.97 | 0.86 |
| Other | 0.99 | 0.98 | 0.99 | 0.90 |

To analyze the model’s class-wise performance and misclassification patterns in greater depth, a confusion matrix was employed (Fig 11). For the Wildboar class, the model correctly classified 900 out of 984 samples, achieving an individual accuracy of 91.46%. The most frequent misclassification occurred with the Deers class (31 cases, 3.15%), likely due to similar body shapes under low-light conditions such as nighttime photography. Misclassifications into Goral (23 cases, 2.34%) and Other (30 cases, 3.05%) were relatively evenly distributed. The Deers class achieved an exceptionally high accuracy of 97.73%, correctly classifying 1,934 out of 1,979 samples. Among 45 misclassification cases, 32 (1.62%) were mistaken for *Other.*, while only a small number were misclassified as Wildboar (8 cases, 0.40%) or Goral (5 cases, 0.25%), indicating that the model effectively learned the unique morphological features of deer. The *Goral* class also achieved a high accuracy of 96.30%, with 1,067 correct predictions out of 1,108 samples. Due to their distinctive morphology, Gorals were relatively easy to distinguish, and misclassifications (41 cases in total) were distributed across Deers (13 cases, 1.17%), Other (17 cases, 1.53%), and Wildboar (11 cases, 0.99%). The *Other* class demonstrated the highest accuracy at 99.61%, with 6,967 correct predictions out of 6,994 samples. This performance can be attributed to the morphological distinctiveness of small and medium-sized mammals (e.g., raccoons, badgers) included in this category, which are clearly distinguishable from the three main species. Misclassifications were extremely rare (27 cases in total), with error rates under 0.2%, confirming the model’s stable performance.

**Figure 11.**
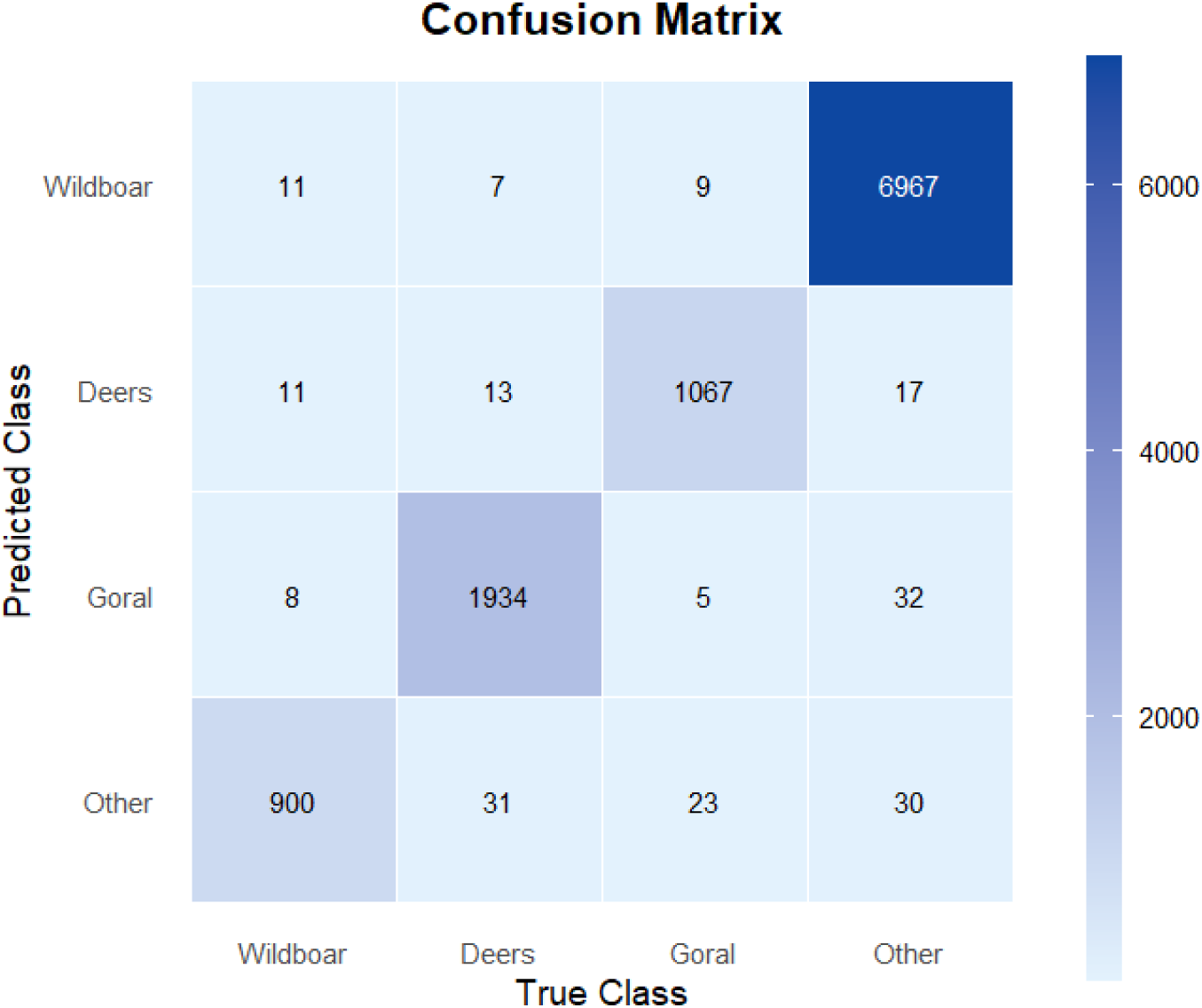
Confusion Matrix of the proposed model. The rows represent the predicted classes, while the columns represent the true (actual) classes. The diagonal elements show the number of correctly classified samples for each class.

Overall, the misclassification analysis revealed that the most frequent error pattern occurred between Wildboar and Deers, reflecting their shared body size and behavioral characteristics as large mammals. By contrast, *Goral* exhibited relatively independent classification performance, indicating strong morphological distinctiveness compared to other classes.

### 4.3. Model Performance Comparison with Baseline Models

To validate the performance of the proposed Graph Attention Contrastive Learning (GACL) model, we conducted comparative experiments against two robust baseline models. The first was YOLOv7, a high-performance object detection model representing a standard, state-of-the-art approach. The second was CACL (Convolutional Attention Contrastive Learning), in which the core Graph Attention module of GACL was replaced with a traditional Convolutional Attention module. All models were trained and evaluated on the same datasets, and their performances were comprehensively analyzed using Precision, Recall, F1-Score, and mAP metrics (Table 4).

**Table 4.** Class-wise performance comparison of the models.

| Model | Metric | Wildboar | Goral | Deers | Others |
| --- | --- | --- | --- | --- | --- |
| YOLOv7 | Precision | 0.58 | 0.95 | 0.65 | 0.82 |
|  | Recall | 0.61 | 0.66 | 0.66 | 0.85 |
|  | **F1-Score** | **0.59** | **0.78** | **0.65** | **0.86** |
| CACL | Precision | 0.93 | 0.75 | 0.96 | 0.89 |
|  | Recall | 0.70 | 0.89 | 0.77 | 0.96 |
|  | **F1-Score** | **0.90** | **0.81** | **0.85** | **0.92** |
| GACL(Proposed) | Precision | 0.91 | 0.96 | 0.97 | 0.99 |
|  | Recall | 0.96 | 0.96 | 0.97 | 0.98 |
|  | **F1-Score** | **0.94** | **0.96** | **0.97** | **0.99** |

The comparative results demonstrate that GACL achieved the most superior and stable performance overall. YOLOv7 yielded relatively low F1-Scores for Wildboar (0.59) and Deers (0.65), indicating limitations in classifying diverse morphological features of Korean wildlife. While CACL showed overall improvement over YOLOv7, issues remained: for Wildboar, CACL achieved higher precision (0.93) but lower recall (0.70), meaning it often missed actual instances. In contrast, GACL attained balanced F1-Scores across all three classes—Wildboar (0.94), Goral (0.96), and Deers (0.97)—highlighting its ability to both identify and correctly classify animals without omission, a capability superior to the baselines.

GACL’s superior performance can be attributed to the synergistic effect of Graph Attention and Contrastive Learning. By explicitly modeling relationships among object parts in a graph structure, GACL captures subtle interactions and contextual cues through its attention mechanism, enabling clear discrimination of morphologically similar species often misclassified by YOLOv7 and CACL. Furthermore, Contrastive Learning reinforces semantic consistency between visual and textual information, maximizing classification accuracy. Conversely, YOLOv7 processes the entire image at once and overlooks inter-object relationships, limiting discrimination in complex backgrounds. While CACL incorporates convolution-based attention to partially model object relationships, it falls short of learning the complex and abstract interactions effectively captured by GACL’s graph-based architecture. This performance gap is consistently reflected in the mAP metric. For the Other class, GACL obtained 0.90, outperforming YOLOv7 (0.83) and CACL (0.89), indicating superior object localization and fine-grained discrimination capabilities.

In conclusion, GACL’s performance advantage stems from the explicit learning of inter-object relationships via Graph Attention and the enhancement of semantic consistency through Contrastive Learning, which together overcome the limitations of YOLOv7 and CACL and enable robust and precise wildlife classification.

### 4.4. Model Performance Comparison with a Global System

To evaluate the practical performance and field applicability of the proposed model, a comparative analysis was conducted against the global platform Wildlife Insights. Wildlife Insights is a large-scale platform that analyzes camera trap images worldwide using Google’s high-performance AI models. To ensure objectivity and reliability, a completely independent validation set was newly constructed, separate from the data used in previous training and testing phases. This validation set, collected from diverse habitats across Korea, comprised 15,000 images for each of the four classes (Wild Boar, Goral, Deers, and Other), along with an additional 15,000 images containing no animals (e.g., empty frames, vehicles, humans). By directly comparing the classification results from our proposed model with those obtained from the Wildlife Insights system on the same dataset, we aimed to quantitatively assess the performance gap between a general-purpose global model and a region-specific model.

The comparative analysis demonstrated that the proposed model exhibited overall higher reliability and accuracy for domestic environmental datasets compared to the global, general-purpose model *Wildlife Insights.* The core performance metrics of the two models are summarized in Table 5, while the complete classification results (Appendix B) and detailed misclassification patterns derived from confusion matrix analyses (Appendix C) are provided in the Appendix.

**Table 5.** Comparative Analysis of Key Performance Metrics

| True Class | Metric | Proposed Model | Global Model<br>(Wildlife Insights) |
| --- | --- | --- | --- |
| Wildboar | Accuracy (Recall) | 90.4% (13,559) | 91.1% (13,667) |
|  | Misclassified as Other | 1.8% (269) | 6.5% (981) |
| Deers | Accuracy (Recall) | 92.4% (13,865) | 82.9% (12,442) |
|  | Misclassified as Other | 1.7% (256) | 15.8% (2,368) |
| Goral | Accuracy (Recall) | 95.4% (14,314) | 33.5% (5,018) |
|  | Misclassified as Other | 1.6% (234) | 60.8% (9,119) |
| ECH | Accuracy (Recall) | 92.5% (13,874) | 90.6% (13,588) |
|  | Misclassified as Other | 7.5% (1,126) | 9.4% (1,412) |

The most significant performance disparity was observed in the Goral (*Naemorhedus caudatus*). The proposed model achieved a high accuracy of 95.4%, whereas the global model correctly classified only 5,018 images (33.5%) out of 15,000 actual Goral images. A total of 9,119 images (60.8%) were misclassified as ‘Other’, among which 4,016 failed to provide any species-level result, while a considerable number were erroneously classified as Deers (601 cases) or Wildboar (246 cases). For the Deers class, while the global model also achieved a relatively high accuracy of 82.9% for the *Cervidae* family, the proposed model demonstrated a more stable performance with an accuracy of 92.4%. For the Wildboar class, both models performed comparably, accurately identifying over 90% at the species level. This is likely attributable to the cosmopolitan distribution of wild boars, which would have allowed a global model like Wildlife Insights to be trained on a vast and diverse dataset for this species. Evaluation of the “Other” class required a cautious approach due to the fundamental differences in class definition between the two models. In the proposed model, ‘Other functions as a single category encompassing all animals except the three predefined classes (Wildboar, Goral, and Deers). In contrast, the global model includes dozens of animal classes from around the world, classifying the same images into much more fine-grained categories. Consequently, rather than a direct comparison of true positive rates, it was more appropriate to assess the models’ limitations from the perspective of false negatives (FN) and false positives (FP). While both models showed a low false positive rate for ’Other’ animals being misclassified into the three main classes, they exhibited a relatively high false negative rate (Proposed: 13.7%, Global: 14.1%), where animals were present in the images but remained undetected. This tendency was even more pronounced in the ECH (Empty, Cars, Human) class, which represents images without animals. For this dataset, the global model correctly identified 13,588 images (90.6%) as empty. However, the remaining 9.4%, amounting to 1,427 images, resulted in false positive errors, frequently misinterpreting background elements such as branches or objects as animals. Such errors introduce a factor that increases the manual workload for researchers during the data cleaning process. In contrast, the proposed model reduced this error rate to 7.5%, demonstrating a tangible improvement in analytical efficiency.

## 5. Conclusion and Discussion

This study was initiated to address the bottleneck in wildlife camera-trap image analysis and, in particular, was motivated by the need for a high-performance automated system tailored to the specific characteristics of the Korean ecological environment. To this end, we constructed a large-scale “Korean Wildlife Dataset” using data directly collected from diverse domestic habitats and developed a novel analysis pipeline that integrates a Graph Attention Transformer (GAT) with Parallel Contrastive Learning. Overall, the proposed model demonstrated highly reliable classification performance across four major wildlife classes, achieving an overall accuracy of 96.83%, which can be regarded as sufficiently applicable to real-world wildlife monitoring systems. However, the true significance of this study lies beyond achieving high accuracy. Its key contribution is the demonstrated advantage of the proposed model in precisely recognizing endemic species, as revealed through comparative analysis with a global general-purpose model. For example, when compared with the global model, its limitations became evident, particularly in fine-grained, species-level analysis. In the case of the long-tailed goral (*Naemorhedus caudatus*), while the global model correctly categorized many images into the broader taxonomic group of *Bovidae*, it identified the species as the long-tailed goral in only 5,018 out of 15,000 cases (33.5%) (Appendix B). This result indicates that although the global model could recognize gorals as generic *Bovidae* animals, it failed to capture the unique species-level features of the native Korean goral. Particularly for the “Other” class, which included diverse species such as raccoon dogs (*Nyctereutes procyonoides*) and badgers (*Meles meles*), correct species-level classification was exceedingly rare, and even family-level identification was often inaccurate. These findings strongly highlight the importance of developing region-specialized models. By being trained intensively on domestic ecological data, our proposed model successfully captured subtle characteristics of local endemic species that global models overlooked, thereby achieving higher classification accuracy. Thus, the significance of this study lies not only in its technical achievement of developing a high-performance AI model but also in its empirical demonstration that, for effective biodiversity conservation, the development of localized AI models that reflect the ecological specificities of the region is not merely beneficial but essential (Tabak et al., 2019).

Of course, the proposed GACL model also has several limitations, which in turn suggest important directions for future research. First, fine-grained classification among morphologically similar large mammals remains challenging, and there is still room for improvement within our model. For example, wild boars and cervids often share similar silhouettes under certain camera angles or low-light conditions, which increases the likelihood of misclassification. To address this limitation, future research could incorporate multi-scale learning to simultaneously consider features at various scales (Lin et al., 2017; Liu et al., 2018; Yang et al., 2016), or utilize multi-modal data, such as thermal imagery, to establish classification criteria that go beyond morphological similarity (Hwang et al., 2015; Kellenberger et al., 2018; Sharma et al., 2025). Second, as with other high-performance models, performance inevitably degrades under extremely low-light conditions or when analyzing low-resolution images (Li et al., 2021; Tang et al., 2023). This is a widely recognized challenge in computer vision, particularly within the context of camera trap image analysis (Yang et al., 2024). Although the GACL model achieved strong overall performance, recognition accuracy can still decline in environments where the extraction of visual features is physically constrained by darkness. Future work should focus on advancing feature extraction algorithms and reinforcing robustness through continuous validation with high-resolution and adverse-condition data (Michaelis et al., 2019). This issue is especially critical in environments like Korea, where seasonal cycles are pronounced and background conditions fluctuate significantly with climate. It is therefore essential to develop models that can maintain robust performance under such seasonal and environmental variations. Enhancing resilience against challenges such as snow-covered landscapes, changes in forest density, or fog is vital for ensuring reliable long-term monitoring (Sakaridis et al., 2021). Finally, a long-term research direction to expand the model’s practical utility lies in lightweighting, with the ultimate goal of enabling real-time processing directly in the field. Since ecological monitoring often takes place in remote mountainous areas under low-power and low-computation constraints, developing lightweight models that can operate stably in such environments would hold immense practical value by providing immediate ecological insights on-site. Through these follow-up studies, the technological foundation established in this research can be further advanced, thereby making a tangible contribution to elevating the scientific capacity of biodiversity conservation and wildlife management in Korea.

## AUTHOR CONTRIBUTIONS

Youngmin Kim conceived the ideas, designed the methodology, collected the data, and performed the analysis. Youngmin Kim, Chul-Han Kim, and Chang-Seop Yun contributed to the analysis, developed the model, and wrote the script. Gea-Jae Joo provided valuable review and discussion throughout the project. Y Kim, CH Kim, and CS led the writing of the manuscript and analyzed the data. All authors contributed critically to the drafts and gave final approval for publication.

## Appendix A. Detailed Data Augmentation Methods

### A.1. Geometric Transformations

**1. Resize:** To achieve scale invariance, images were randomly resized using both upscaling (magnification factors of ×2, ×4, ×6, ×8) and downscaling (reduction factors of 1/2, 1/4, 1/6, 1/8). Bilinear interpolation was employed as the interpolation method. The transformation is formulated as:

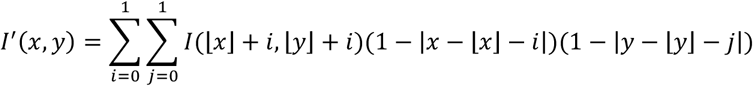

where *I*(*x*, *y*) denotes the pixel value of the original image and, *I*^′^(*x*, *y*) denotes the pixel value in the transformed image.

**2. Rotation:** To achieve rotation invariance, images were rotated by an angle *θ* randomly selected from [-360°, 360°]. The transformation was implemented using a 2D rotation matrix. Any empty regions created post-rotation were handled using nearest-neighbor interpolation. The rotation matrix is defined as:

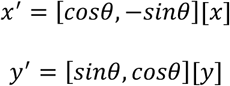

where *θ* represents the rotation angle (−360^°^ ≤ *θ* ≤ 360^°^). Any empty regions created during rotation were filled using nearest-neighbor interpolation

### A.2. Photometric Transformations

To simulate diverse lighting and color conditions, a dual-space approach leveraging both the HSV and LAB color spaces was adopted.

**3. HSV Color Space:** Adjustments were made to saturation, exposure, and hue with equal probability.

**Saturation (S)** was adjusted by

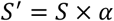

where *S* is the original saturation value and *α* is the adjustment factor.

**Exposure (L)** was implemented via gamma correction: L’ = L^(1/γ), where γ ranges from 0.1 to 10.0.

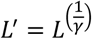

where *γ* denotes the gamma value(0.1 ≤ *γ* ≤ 10.0).

**Hue (H)** was adjusted by a cyclical shift within the range

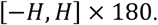

These three transformations—saturation, exposure, and hue adjustments—were applied with equal probability (1:1:1).

**4. LAB Color Space:** To achieve more perceptually uniform variations, adjustments were made to the a* (green-red) and b* (blue-yellow) channels:

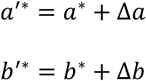

where the adjustment factors Δ*a* and Δ*b* are each selected from a range of -50 to +50.

To quantify the increased diversity in color distribution, the Shannon entropy metric was employed. The information entropy for each channel’s histogram is defined as:

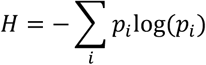

where *p*_*i*_ is the normalized probability of the *i*-th bin.

A comparative analysis of the two color spaces was conducted using the Kullback-Leibler (KL) divergence.

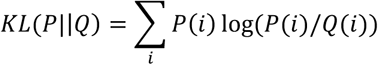

### A.3. Mixup Augmentation

Mixup was used for regularization. New samples (*x*~, *y*~) were defined as:

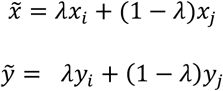

where *i* and *j* are two distinct training samples, and the mixing ratio *λ* is randomly sampled from a Beta distribution.

## Appendix B. Detailed Species-Level Classification Results of the Global Model (Wildlife Insights)

This table presents the detailed classification results of the global model on the validation set. Each sub-table corresponds to one of the five true (actual) classes. The ’Predicted Species’ column lists the common names assigned by the model, and the ’Count’ column shows the number of images assigned to each prediction.

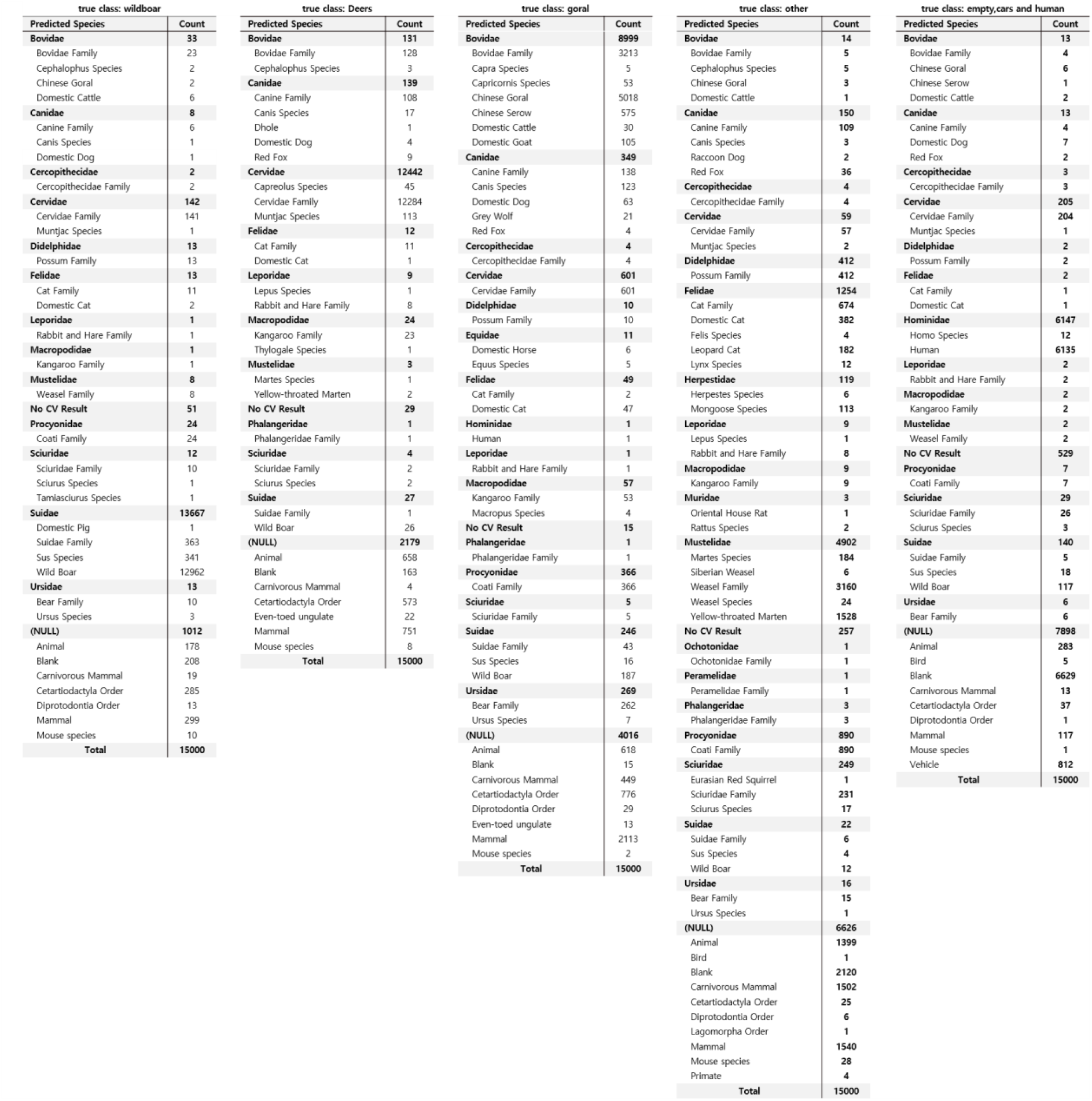

## Appendix C. Comparative Analysis of Confusion Matrices

A side-by-side comparison of the confusion matrices for the **proposed model (left)** and the **global model (right)** on the validation set. The rows represent the predicted class, and the columns represent the true (actual) class.

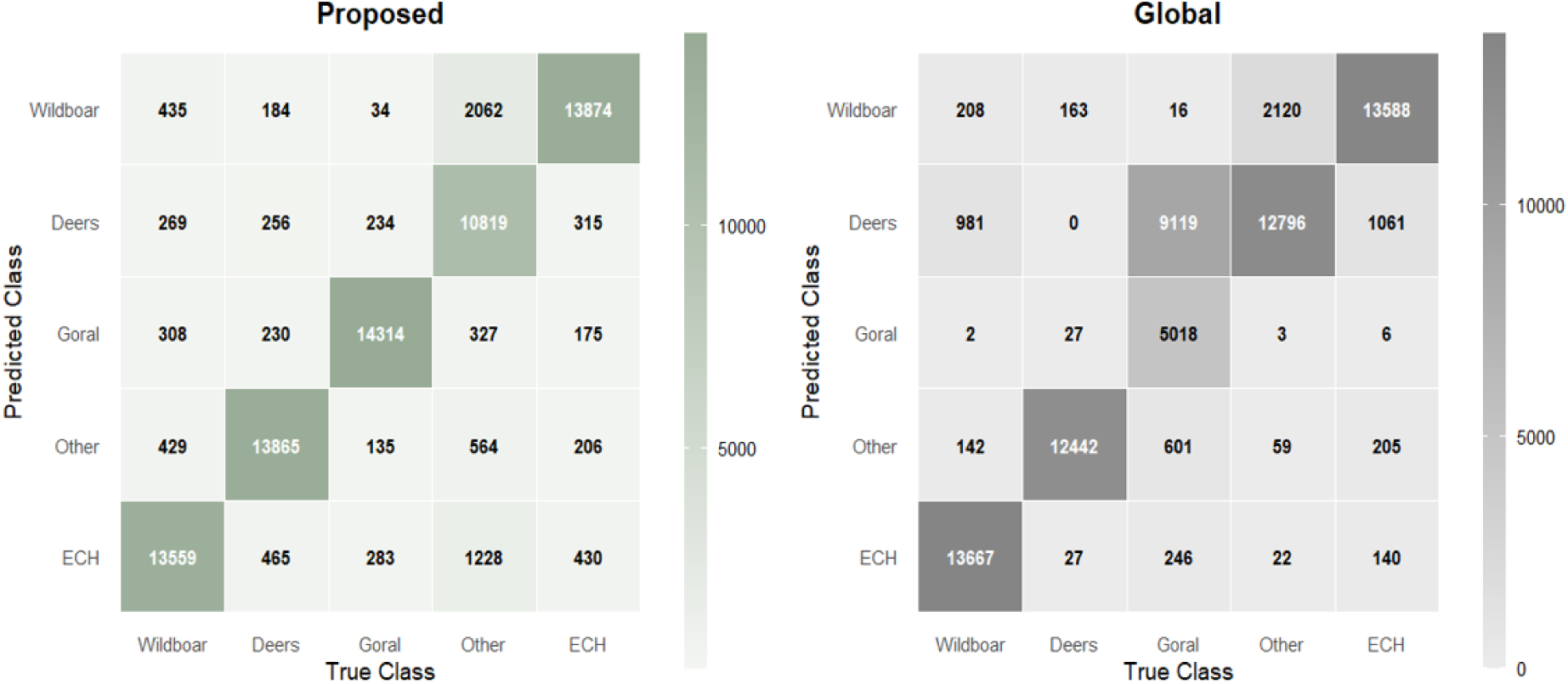

## References

1. Ahumada, J. A., Hurtado, J., & Lizcano, D. (2013). Monitoring the status and trends of tropical forest terrestrial vertebrate communities from camera trap data: A tool for conservation. PLoS ONE, 8(9), e73707.

2. Ahumada, J. A., Silva, C. E., Gajapersad, K., Hallam, C., Hurtado, J., Martin, E., McWilliam, A., Mugerwa, B., O’Brien, T., Price, J., & Rovero, F. (2011). Community structure and diversity of tropical forest mammals: data from a global camera trap network. Philosophical Transactions of the Royal Society B: Biological Sciences, 366(1578), 2703–2711.

3. Beery, S., Morris, D., & Yang, S. (2019). Efficient pipeline for camera trap image review. arXiv preprint arXiv:1907.06772.

4. Cai, Z., Fan, Q., Feris, R., & Vasconcelos, N. (2016). A unified multi-scale deep convolutional neural network for fast object detection. In European Conference on Computer Vision. Cham: Springer International Publishing.

5. Fennell, M., Beirne, C., & Burton, A. C. (2022). Use of object detection in camera trap image identification: Assessing a method to rapidly and accurately classify human and animal detections for research and application in recreation ecology. Global Ecology and Conservation, 35, e02104.

6. Hernández-García, A., & König, P. (2018). Further advantages of data augmentation on convolutional neural networks. In International Conference on Artificial Neural Networks (pp. 95– 104). Cham: Springer International Publishing.

7. Hwang, S., Park, J., Kim, N., Choi, Y., & So Kweon, I. (2015). Multispectral pedestrian detection: Benchmark dataset and baseline. In Proceedings of the IEEE Conference on Computer Vision and Pattern Recognition.

8. Ji, Y., Zhang, H., & Wu, Q. M. J. (2018). Salient object detection via multi-scale attention CNN. Neurocomputing, 322, 130–140.

9. Kellenberger, B., Marcos, D., & Tuia, D. (2018). Detecting mammals in UAV images: Best practices to address a substantially imbalanced dataset with deep learning. Remote Sensing of Environment, 216, 139–153.

10. Krizhevsky, A., Sutskever, I., & Hinton, G. E. (2017). ImageNet classification with deep convolutional neural networks. Communications of the ACM, 60(6), 84–90.

11. Li, C., Guo, C., Han, L., Jiang, J., Cheng, M. M., Gu, J., & Loy, C. C. (2021). Low-light image and video enhancement using deep learning: A survey. IEEE Transactions on Pattern Analysis and Machine Intelligence, 44(12), 9396–9416.

12. Lin, T. Y., Goyal, P., Girshick, R., He, K., & Dollár, P. (2017). Feature pyramid networks for object detection. In Proceedings of the IEEE Conference on Computer Vision and Pattern Recognition.

13. Ma, Z., Jin, X., Pan, X., & Xu, Z. (2024). Wildlife real-time detection in complex forest scenes based on YOLOv5s deep learning network. Remote Sensing, 16(8), 1350.

14. Mathis, A., Mamidanna, P., Cury, K. M., Abe, T., Lo Sátyro, S., Lindsay, G. W., … & Mathis, M. W. (2018). DeepLabCut: markerless pose estimation of user-defined body parts with deep learning. Nature Neuroscience, 21(9), 1281–1289.

15. Michaelis, C., Mitzkus, B., Geirhos, R., Rusak, E., Bringmann, O., Ecker, A. S., Kohl, T., & Brendel, W. (2019). Benchmarking robustness in object detection: Autonomous driving when winter is coming. arXiv preprint arXiv:1907.07484.

16. Norouzzadeh, M. S., Nguyen, A., Kosmala, M., Swanson, A., Palmer, M. S., Packer, C., & Clune, J. (2018). Automatically identifying, counting, and describing wild animals in camera-trap images with deep learning. Proceedings of the National Academy of Sciences, 115(25), E5716– E5725.

17. Raj, S., Kumari, S., Kakarla, A., & Verma, M. (2025). Deep Learning-Based Object Detection in Thermal Imaging. 2025 Global Conference in Emerging Technology (GINOTECH). IEEE.

18. Sakaridis, C., Dai, D., & Van Gool, L. (2021). ACDC: The adverse conditions dataset with correspondences for semantic driving scene understanding. In Proceedings of the IEEE/CVF International Conference on Computer Vision (pp. 10765–10775).

19. Tabak, M. A., Norouzzadeh, M. S., Wolfson, D. W., Newton, E., Chandler, J. I., Plumer, D., … & Miller, R. S. (2019). Machine learning to classify animal species in camera trap images: Applications in ecology. Methods in Ecology and Evolution, 10(4), 585–590.

20. Tang, H., Zhu, H., Fei, L., Wang, T., Cao, Y., & Xie, C. (2023). Low-illumination image enhancement based on deep learning techniques: A brief review. Photonics, 10(2), 198.

21. Wei, X. S., Song, Z., Wu, Y., & Sun, J. (2021). Fine-grained image analysis with deep learning: A survey. IEEE Transactions on Pattern Analysis and Machine Intelligence, 44(12), 8927–8948.

22. Wildlife Insights (2020). Wildlife Insights: A platform for wildlife monitoring. https://www.wildlifeinsights.org

23. Willi, M., Pitman, R. T., Cardoso, A. W., Locke, C., Swanson, A., Boyer, A., … & Fortson, L. (2019). Identifying animal species in camera trap images using deep learning and citizen science. Methods in Ecology and Evolution, 10(1), 80–91.

24. Yang, Z., Tian, Y., & Zhang, J. (2024). Adaptive image processing embedding to make the ecological tasks of deep learning more robust on camera traps images. Ecological Informatics, 82, 102705.

25. Zhang, H., Cisse, M., Dauphin, Y. N., & Lopez-Paz, D. (2017). mixup: Beyond empirical risk minimization. arXiv preprint arXiv:1710.09412.

26. Zou, Z., Shi, Z., Guo, Y., & Ye, X. (2023). Object detection in 20 years: A survey. Proceedings of the IEEE, 111(3), 257–276.

